# Translocator Protein Localises To CD11b^+^ Macrophages In Atherosclerosis

**DOI:** 10.1101/494377

**Authors:** Chantal Kopecky, Elvis Pandzic, Arvind Parmar, Jeremy Szajer, Victoria Lee, Alexander Dupuy, Andrew Arthur, Sandra Fok, Renee Whan, William J. Ryder, Kerry-Anne Rye, Blake J. Cochran

## Abstract

**Background:** Atherosclerosis is characterized by lipid deposition, monocyte infiltration and foam cell formation in the artery wall. Translocator protein (TSPO) is abundantly expressed in lipid rich tissues. Recently, TSPO has been identified as a potential diagnostic tool in cardiovascular disease. The purpose of this study was to determine if the TSPO ligand, ^18^F-PBR111, can identify early atherosclerotic lesions and if TSPO expression can be used to identify distinct macrophage populations during lesion progression.

**Methods and Results:** ApoE^−/−^ mice were maintained on a high-fat diet for 3 or 12 weeks. C57BL/6J mice maintained on chow diet served as controls. Mice were administered ^18^F-PBR111 intravenously and PET/CT imaged. After euthanasia, aortas were isolated, fixed and optically cleared. Cleared aortas were immunostained with DAPI, and fluorescently labelled with antibodies to-TSPO, the tissue resident macrophage marker F4/80 and the monocyte-derived macrophage marker CD11b. TSPO expression and the macrophage markers were visualised in fatty streaks and mature lesions by light sheet microscopy. While tissue resident F4/80^+^ macrophages were evident in the arteries of animals without atherosclerosis, no CD11b^+^ macrophages were observed in these animals. In contrast, mature plaques had high CD11b and low F4/80 expression. A ~3-fold increase in the uptake of ^18^F-PBR111 was observed in the aortas of atherosclerotic mice relative to controls.

**Conclusions:** Imaging of TSPO expression is a new approach for studying atherosclerotic lesion progression and inflammatory cell infiltration. The TSPO ligand, ^18^F-PBR111, is a potential clinical diagnostic tool for the detection and quantification of atherosclerotic lesion progression in humans.

## Introduction

Atherosclerosis, a chronic inflammatory disease characterized by accumulation of lipids and inflammatory cells such as macrophage foam cells within the arterial wall, is a leading cause of death globally^1^. There is significant debate as to the relative contributions of tissue-resident versus monocyte-derived macrophages to atherosclerotic lesion progression^2^.

Atherosclerotic progression commonly develops as an indolent process over decades^3^, making early identification of the condition challenging. Clinical assessment of atherosclerosis currently involves either invasive angiography or intravascular ultrasound (IVUS), or is limited to recognition of advanced disease by, for example, calcium scoring^4^. Recent efforts have focused on establishing dynamic imaging approaches to identify clinically relevant atherosclerosis. The development, assessment and broad adoption of novel PET/CT techniques enables early detection and diagnosis of cardiovascular disease^5^. Non-invasive PET imaging, using specifically targeted radiolabelled probes to track active atherosclerosis may provide a highly suitable method to identify and further understand atherogenesis^6^. In this regard, translocator protein (TSPO) has recently been identified as a therapeutic target and diagnostic tool in cardiovascular disease^7^.

TSPO is an 18 kDa mitochondrial membrane protein that is involved in steroidogenesis and cholesterol transport^8^. It is expressed abundantly throughout the body, and is highly upregulated in microglia and macrophages during inflammation^9,10^. Clinically, the TSPO radioligand ^11^C-PK11195 has been used to image atherosclerotic lesions in the carotid artery^11^. However, use of this ligand was limited due to uptake by healthy vessels. Newer TSPO ligands have been developed in recent years and ^18^F-PBR111 has evolved as promising candidate owing to its higher affinity and binding specificity compared to ^11^C-PK11195^12^. Moreover, as TSPO is involved in the regulation of cardiovascular, atherosclerotic and inflammatory processes, TSPO ligands are promising therapeutic and diagnostic tools for use in cardiovascular disease^7^. Treating macrophages with TSPO agonists has been shown to increase cholesterol efflux^13^ and treatment of cholesterol fed guinea pigs and apoE^−/−^ mice with a peptide containing the cholesterol binding domain of TSPO reduced serum cholesterol levels and decreased atheroma formation^14^.

In this study, the ability of ^18^F-PBR111 to detect atherosclerosis progression *in vivo* was examined. Specifically, the ability of ^18^F-PBR111 to detect atherosclerotic lesions at different stages of development was investigated. It also asked if TSPO expression increases with disease progression and if distinct macrophage populations can be distinguished by TSPO expression during lesion development. The findings of this study provide a rationale for the development of TSPO radioligands as clinical diagnostic tools for the detection and quantification of atherosclerosis in humans by non-invasive, biomedical imaging.

## Methods

### Mice

C57BL/6J and B6.129P2-*Apoe^tm1Unc^* (apoE^−/−^) mice were obtained from Animal Resources Centre (Perth, Australia) and maintained in pathogen-free conditions at the Brain and Mind Centre, University of Sydney. Control C57BL/6J mice were fed a standard chow diet, whilst apoE^−/−^ mice were maintained on a high fat diet (HFD; 22% fat, 0.15% cholesterol, Specialty Feeds, Perth, Australia) for the time periods stated. All procedures were approved by the University of Sydney Animal Ethics Committee and followed the NIH *Guide for the care and use of laboratory animals*, Eighth edition (2011).

### Light sheet microscopy

Mice were euthanized after PET/CT scanning and aortas were isolated and fixed overnight in 4% (v/v) paraformaldehyde. Lipid was cleared from the tissue as previously described^15^. Briefly, fixed tissues were incubated in Reagent 1 (25% (w/w) urea, 25% (w/w) Quadrol, 15% (w/w), Triton X-100) for 16 h at 37°C and then washed for 36 h in PBS. Optically cleared samples were then stained with Alexa 488 labelled anti-TSPO antibody (1:500, Abcam), Alexa 647 labelled anti-F4/80 antibody (1:200, Abcam), PE/Dazzle 594 labelled anti-CD11b antibody (1:200, Biolegend) and DAPI (ThermoFisher) in PBS plus 0.1% (w/v) BSA, 0.3% (v/v) Triton X-100 for 16 h at 37 °C. Samples were washed for 36 h in PBS and incubated in refractive index matched Reagent 2 (25% (w/w) urea, 50% (w/w) sucrose, 10% (w/w) triethanolamine) for 2 days. Stained tissues were imaged by lightsheet fluorescence microscopy (Zeiss Lightsheet Z.1; Carl Zeiss) housed at UNSW Sydney (Biomedical Imaging Facility, Mark Wainwright Analytical Centre, UNSW Sydney). The samples were placed in the imaging chamber filled with Reagent 2 and imaging was performed using a 5×/0.16 detection lens and 5×/0.1 illumination lens. The four fluorescent signatures were collected as follows: DAPI 405 nm excitation, 460–500 nm emission; TSPO 488 nm excitation, 505–545 nm emission; CD11b 561 nm excitation, 575–615 nm emission and F4/80 638 nm excitation, 660/LP nm emission. To capture the entire sample, a Z stack of ~1000 frames and ~3×4 tiles were captured, at 30 ms exposure on two CMOS cameras with 1920×1920 pixel images and 2.329 μm pixel size. Z-stack images were collected with a mean optimal sectioning step of 4 μm. Images were processed, stitched, rendered and visualized in 4D with Arivis Vision4D software (Arivis).

### Data quantification and image analysis

Stitched data was exported to tif files for further processing in Matlab 2018a (Mathworks). When imported, data from each channel was registered to align with the DAPI channel in order to correct potential drift artefacts from imaging. Following alignment, TSPO channel data was imported into Microscopy Image Browser^16^ running in Matlab for segmentation of TSPO enriched regions and the aortic wall in each optical section. TSPO 3D masks were then assessed for TSPO^+^ volume, while aorta wall masks were used to calculate total aorta and aortic wall volumes. Using these masks, the standard deviation of TSPO channel intensity within the aorta wall was determined, and the ratio of standard deviation and average intensity calculated for each slice. In order to characterize the amount of cross correlation between TSPO and CD11b, image cross correlation spectroscopy was used^17^. The TSPO^+^ and aorta wall masks described above were applied to the TSPO and CD11b channels in each slice to exclude pixels outside of the mask from the cross correlation function. The cross correlation function was characterized for each slice by fitting to a 2D Gaussian function and the amplitude extracted, with higher amplitude indicating higher cross-correlation between TSPO and CD11b.

### PET study

Mice were maintained on chow or high fat diet as indicated. Mice were anaesthetised with isoflurane administered via precision vaporiser (induction, 4%; maintenance, 1.5–2%), and maintained in a surgical plane of anaesthesia for the duration of the procedure. After positioning the mice in the gantry of an Inveon PET/CT Scanner (Siemens, Munich, Germany), a 60 min PET scan commenced immediately after injection of ^18^F-PBR111 (8–18 MBq, 100 μL, 0.2 nM) via the lateral tail vein. To allow for co-registration of radiotracer uptake with the aorta, a CT scan was performed immediately after the PET scan.

### Image reconstruction

Image reconstruction was performed using IAW 2.02 (Siemens). The listmode data was histogrammed into 16 frames (6 10 s, 4 60 s, 1×300 s, 5×600 s) for the period 0–60 min after tracer injection. Emission sinograms were reconstructed using 2D-filter back projection with a zoom 1.5. The reconstructed images consisted of 16 frames of a 128×128×159 matrix with a voxel size of 0.52 0.52 0.796 mm^3^, corrected for attenuation (CT-based), scatter, randoms, normalisation, isotope decay, branching ratio, deadtime and were calibrated to Bq/mL.

### Region of interest

The thoracic aorta, from the commencement of the descending aorta to the aortic hiatus of the diaphragm, was used as the relevant region of interest (ROI) for quantification. The thoracic aorta rather than the aortic arch was chosen because it did not require cardiac gating. The ROI for each mouse was demarcated using a voxel size of 0.2×0.2×0.2 mm^3^ by a physician trained in medical imaging interpretation. ROIs were delineated using the CT images viewed using AMIDE software^18^ on a 4-megapixel EIZO monitor at the Department of Nuclear Medicine, Concord Hospital, NSW. The interpreting physician was blinded to the identity of each mouse. Each ROI was independently reviewed by two additional interpreters to ensure accurate demarcation of the thoracic aorta. Once the ROIs were agreed upon, the PET images where used to demarcate a secondary ROI within the lateral tail vein, at the base of the tail. This second ROI represented the level of ^18^F-PBR111 in the blood, rostral to the radiopharmaceutical injection site.

### Statistical analysis

All data presented are mean ± SD. Data was analysed using Prism 7 (GraphPad). One-way ANOVA or Kruskal-Wallis tests, as appropriate, were used to determine statistically significant differences between data sets. A *p* value < 0.05 was considered statistically significant.

## Results

### TSPO accumulates in atherosclerotic lesions

Aortas were rendered optically clear (Fig. 1A) and stained with fluorescently labelled TSPO and DAPI antibodies. Expression of TSPO during atherosclerotic lesion development was determined in the cleared and immunostained aortas by 3D imaging using lightsheet fluorescence microscopy. TSPO expression was not observed in the aortas of control mice (Fig. 1B), and was restricted to the vessel wall of aortas of 3-week HFD fed apoE^−/−^ mice (Fig. 1C). 12-week HFD fed apoE^−/−^ mice showed high TSPO expression in both regions of mature plaque and fatty streaks lining the vessel wall (Fig. 1D). Quantification of TSPO+ volume demonstrated a significant increase in 3-week HFD fed apoE^−/−^ mice compared to controls (0.64±0.23% vs 0.04±0.06% TSPO^+^ volume for 3-week HFD and control, respectively; p<0.05). 12 weeks of HFD led to a further significant increase in TSPO+ volume (34.05±24.26% TSPO^+^ volume; p<0.05 vs both control and 3-week groups) (Fig. 1E).

**Figure 1:**
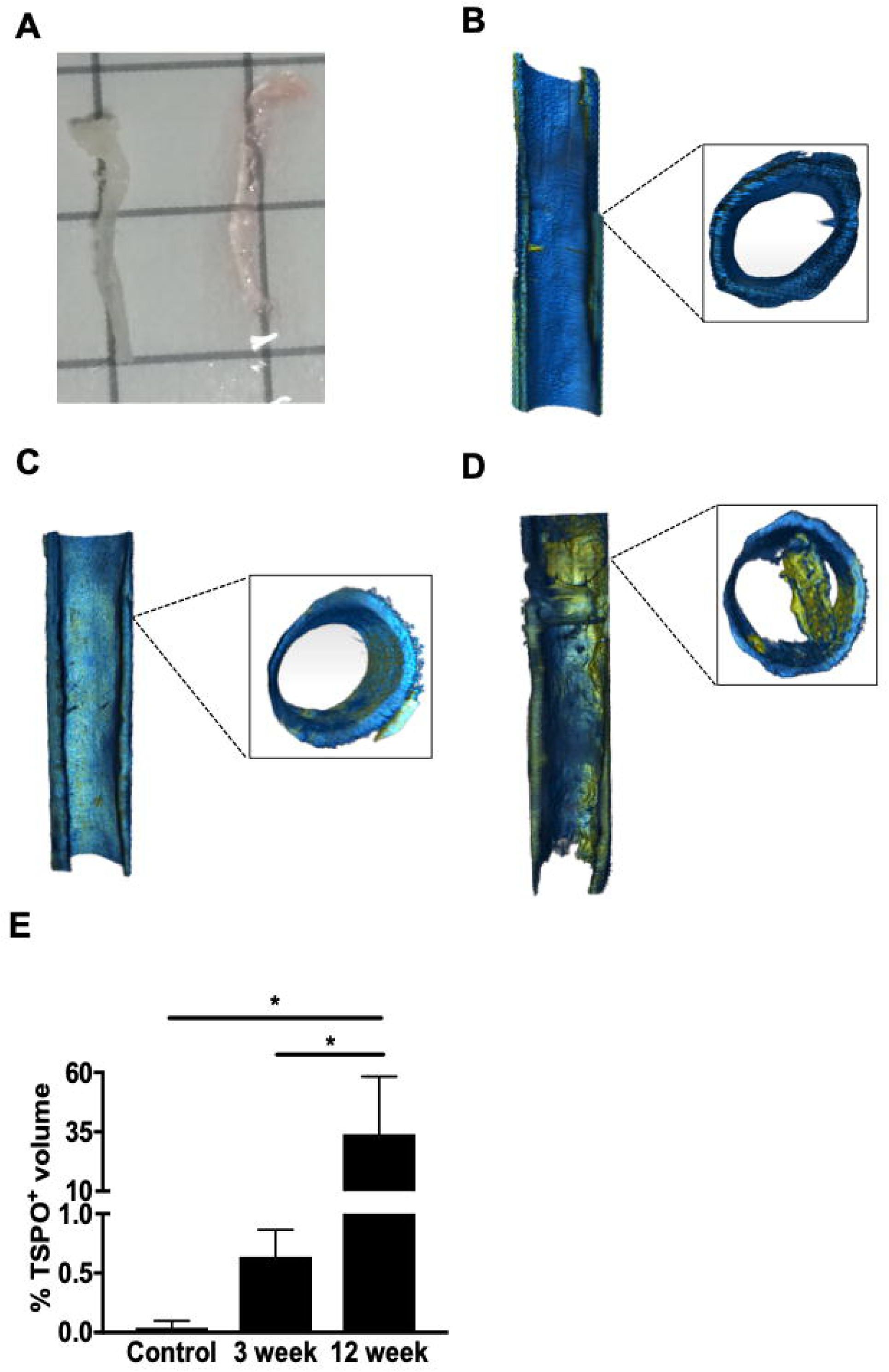
TSPO expression increases with atherosclerosis progression. (**A**) Isolated aorta before (left) and after (right) optical clearing; Representative 3D reconstructed images of aortas isolated from (**B**) C57BL/6J or apoE^−/−^ mice after (**C**) 3- and (**D**) 12-weeks on HFD. TSPO expression is indicated in yellow, DAPI is indicated in blue. (**E**) Quantification of TSPO+ volume in aortas. Values are mean ± SD. n = 4/ group, *p<0.05. Grid in (A) = 1 cm^2^.

To better characterise the pattern of TSPO expression in the aortas of atherosclerotic mice, a masking algorithm was applied (Fig. 2A) to determine a pixel-based intensiometric analysis on a per plane basis, and then calculated the standard deviation (SD) of the TSPO channel intensity within this pre-defined region of interest. These results show that the SD of the TSPO signal in the 12-week HFD fed apoE^−/−^ mice was significantly increased relative to both control and 3-week HFD mice (p<0.05 vs both) (Fig. 2B).

**Figure 2:**
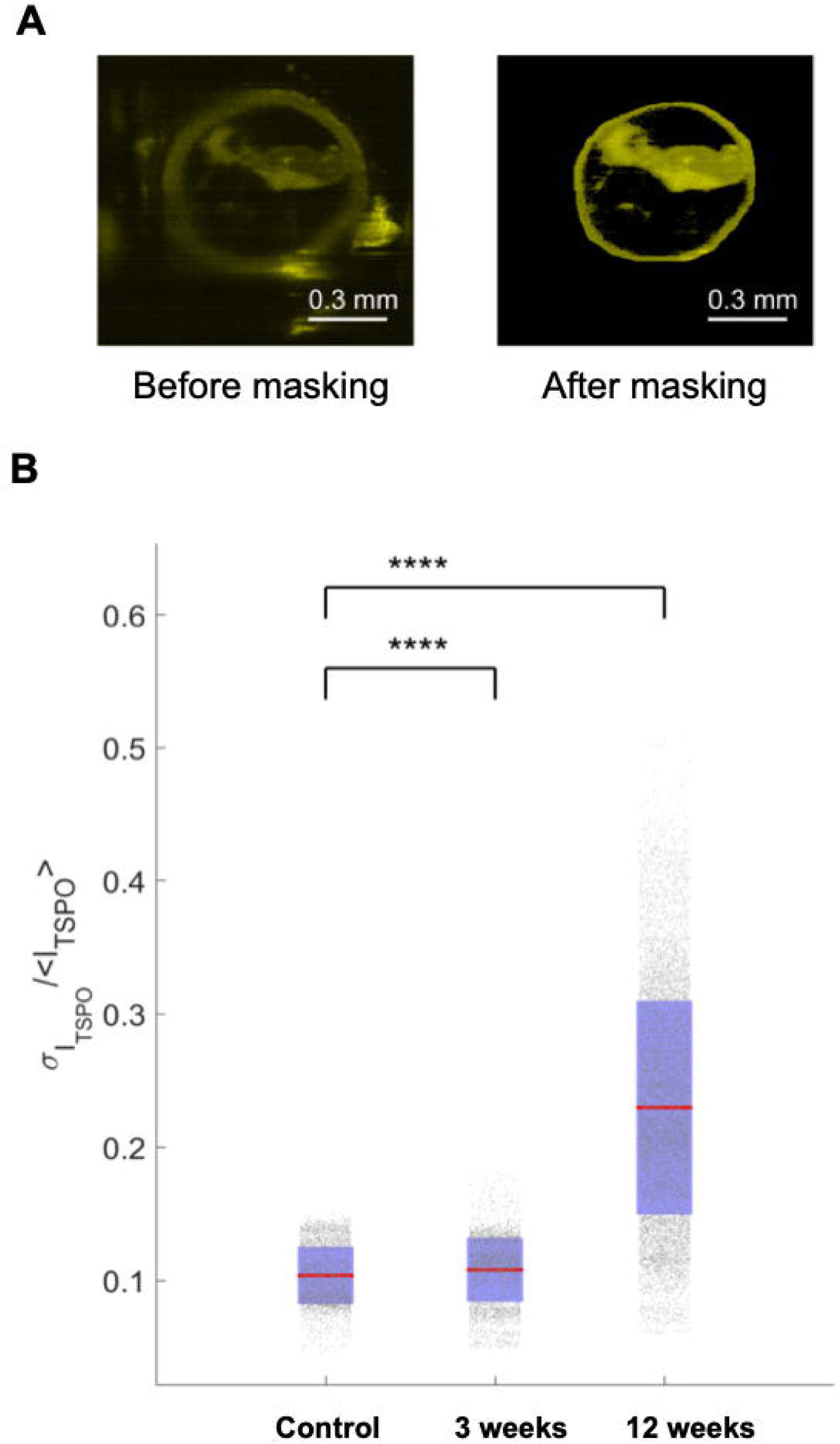
Quantification of the variability of intravascular TSPO expression. (**A**) Representative images before and after a masking algorithm was applied to generate a region of interest and measure TSPO channel intensity within the mask. (**B**) SD of TSPO signal within the mask normalized to TSPO signal in aorta wall. Each dot represents the mean TSPO intensity per plane normalized to the mean overall TSPO intensity of each sample. Red bars indicate means and blue regions indicate confidence intervals. ****p<0.001

### TSPO is preferentially co-localized to CD11b+ macrophages

To investigate the presence of specific macrophage subpopulations throughout atherosclerotic lesion development, aortas were immunostained with the macrophage markers CD11b and F4/80. In aortas from control C57BL/6J mice (Fig. 3A) and 3-week HFD fed apoE^−/−^ mice (Fig. 3B), F4/80^+^ macrophages predominated, while CD11b staining was minimal. In contrast, significant CD11b staining was observed in the aortas of 12-week HFD fed apoE^−/−^ mice (Fig. 3C). Co-localization of the respective macrophage markers and TSPO was determined by calculating the spatial image cross-correlation (ICCS) and expressing the results as the change in ICCS (TSPO/CD11b – TSPO/F4/80) (Figure 3D). High ICCS values, which are indicative of a high correlation between CD11b and TSPO levels, were observed in mature plaque (Fig. 3E) and fatty streak (Fig. 3F) regions. TSPO expression was associated with CD11b expression, as evidenced by the positive ICCS values observed in the vast majority of slices.

**Figure 3:**
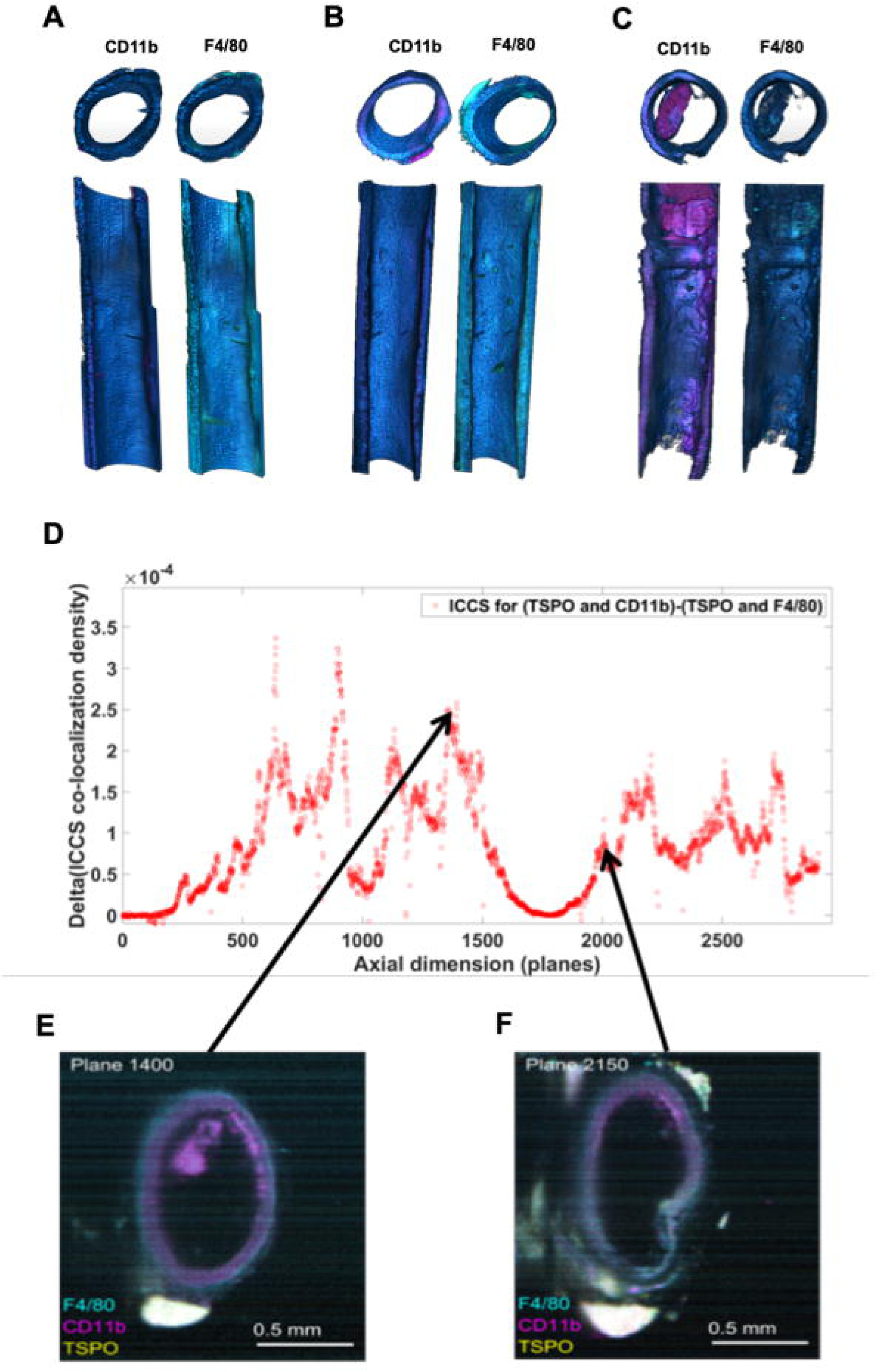
TSPO expression correlates with CD11b. Expression of the macrophage markers CD11b and F4/80 in aortas from (**A**) chow fed wildtype controls (**B**) 3-week HFD and (**C**) 12-week HFD fed apoE^−/−^ mice. (**D**) To quantify co-localization between the macrophage markers CD11b and F4/80 with TSPO, spatial image cross-correlations (ICCS) were calculated and expressed as delta ICCS (TSPO/CD11b – TSPO/F4/80) for each imaging plane. Inserts show expression of markers in representative cross sections of (**E**) mature plaque and (**F**) fatty streak regions.

### ^18^F-PBR111 detects mature, but not early lesions *in vivo*

Given the level of expression and localisation of TSPO in the aortas of apoE^−/−^ mice maintained on HFD, it was next asked if the TSPO agonist ^18^F-PBR111 could also detect atherosclerosis in these mice *in vivo*. To control for the level of circulating ^18^F-PBR111, activity in the lateral tail vein was subtracted from the region of interest in the aorta (Fig. 4). This allowed for the tissue specific uptake in the aorta to be measured. Uptake of ^18^F-PBR111 in the aorta was significantly increased in 12-week high fat fed apoE^−/−^ mice compared to both 3-week HFD apoE^−/−^ and wildtype control mice (maximal activity at 2.7 min post injection; 9.15±3.60 vs 0.77±1.32 and 1.22±1.20 %ID/cm^3^, for 12-week HFD, 3-week HFD and control mice, respectively; p<0.05) (Fig. 5A). Aortic specific levels of ^18^F-PBR111 were significantly increased in 12-week HFD apoE^−/−^ mice over the course of data acquisition (incremental area under the curve: 4015±2808 vs 90±509 and 443±474 for 12-week HFD, 3-week HFD and control mice, respectively; p<0.05 for 12-week HFD vs both) (Fig. 5B).

**Figure 4:**
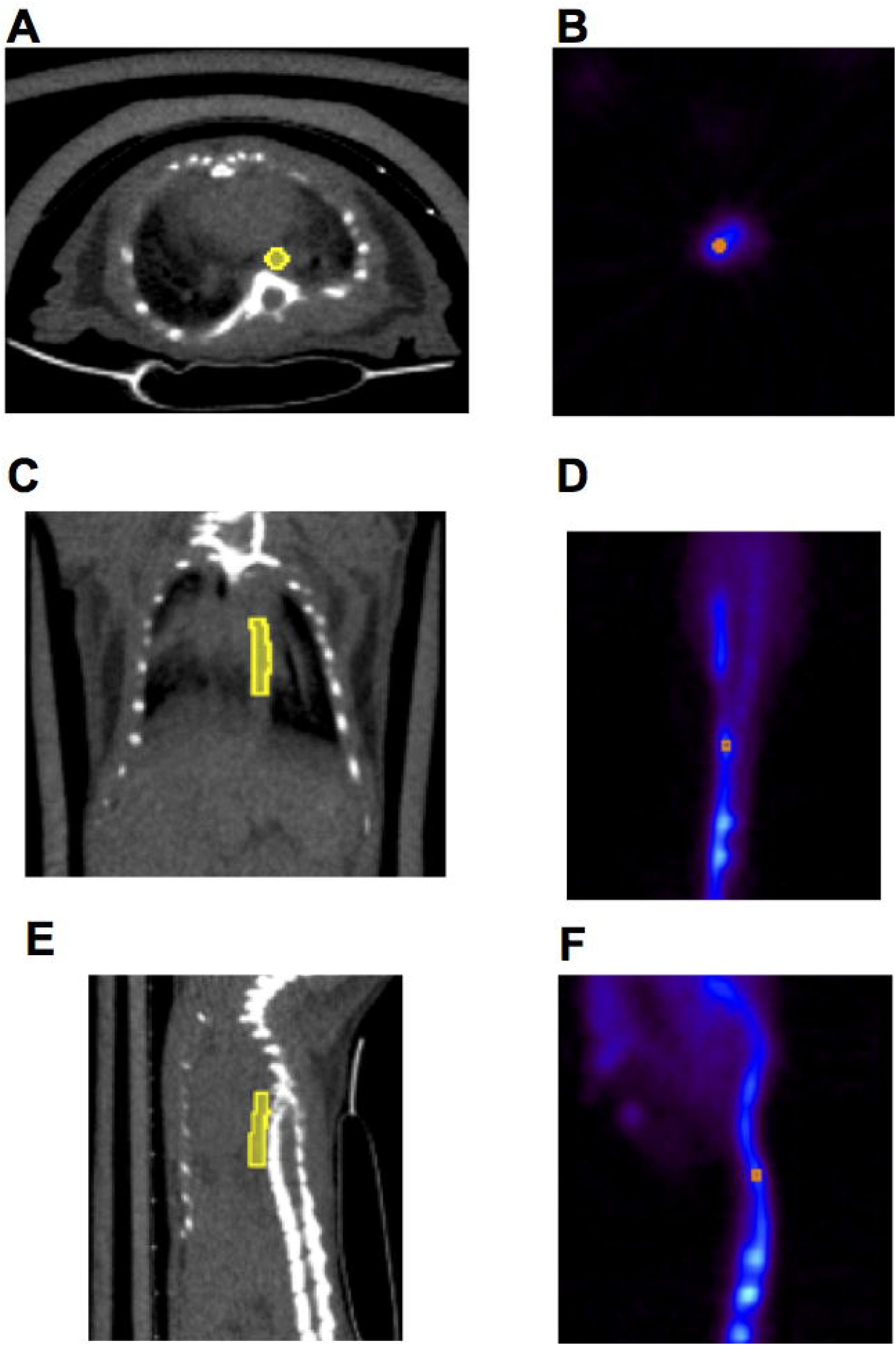
Regions of Interest (ROIs) used in PET imaging of ^18^F-PBR111 aortic uptake. Axial (**A, B**), coronal (**C, D**) and sagittal (**E, F**) views of CT (**A**, **C** and **E**) and PET (**B**, **D** and **F**) images. The aortic ROI is shown in yellow and the lateral tail vein ROI is shown in red.

**Figure 5:**
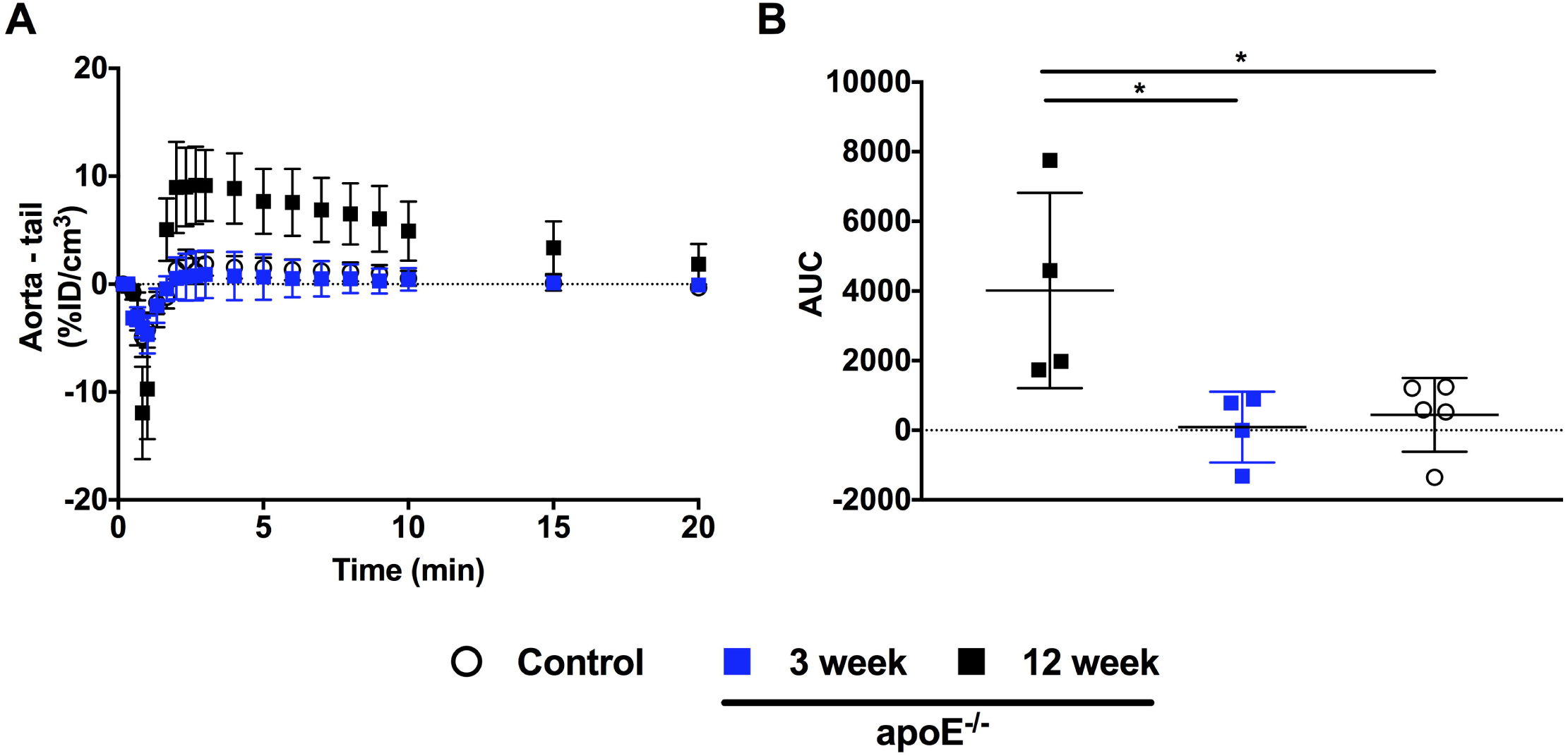
Quantification of ^18^F-PBR111 uptake in thoracic aorta. (**A**) Time-activity curves calculated by subtracting the lateral tail vein ROI from the thoracic aorta ROI and (**B**) corresponding area under curve values. Values are mean ± SD and expressed as percentage of injected dose per cubic cm (ID/cm^3^). n = 4–5/ group, *p<0.05.

## Discussion

This study found that TSPO expression in aortas of high-fat fed apoE^−/−^ mice was progressively elevated with atherosclerotic lesion progression. These results were validated by PET/CT with increased uptake of the TSPO radioligand ^18^F-PBR111 in mice with advanced atherosclerosis relative to mice with early stage disease and control mice without disease. It was also demonstrated that macrophage subpopulations differed between early atherosclerosis and more mature plaque regions. Whilst F4/80+ tissue resident macrophages were evident in aortas of animals without atherosclerosis, no CD11b staining was observed in the arteries of these mice. In contrast, atherosclerotic lesions had significantly increased CD11b expression that co-localized with TSPO.

Atherosclerosis is a leading cause of death globally and is now widely recognized as a lipid-driven, systemic inflammatory disease^19^. A critical initiating event in atherosclerosis is excessive lipid accumulation and foam cell formation within the artery wall, leading to activation of inflammatory signaling^20^. Macrophages and monocytes are the major immune cells in atherosclerosis and play important roles throughout all stages of lesion progression^21^. However, macrophage development is a highly heterogeneous process, and embryonically derived tissue resident macrophages can be joined or replaced by recruited monocytes upon inflammatory stimuli^19, 22^.

This study has shown that CD11b^+^ macrophages predominate in advanced atherosclerotic lesions. Contrastingly, no CD11b staining was present and F4/80^+^ regions were evenly distributed throughout the aortas of control C57BL/6J mice and early lesions in 3 weeks HFD-fed apoE^−/−^ mice. Thus, this data shows for the first time that TSPO accumulates in the same regions of atherosclerotic lesions as specific macrophage subpopulations. There is significant debate regarding the utility and specificity of markers to determine the lineage and source of the macrophages *in vivo^23^*. CD11b is expressed by newly migrating macrophages in the liver^24, 25^, whilst F4/80 is present on the surface of tissue resident macrophages in the spleen and liver^24^.

A challenge in clinical management is that most patients with atherosclerosis do not display overt disease-related symptoms. This also presents a substantial diagnostic challenge^26^. Recent developments have focused on a shift to novel, non-invasive techniques such as PET, coupled with CT or MRI to allow precise anatomical localization and characterisation of atherosclerotic lesions^6^. In agreement with previous studies, the results presented here demonstrate that TSPO may be a useful marker of atherosclerotic lesion development. Furthermore, the specific association of TSPO with CD11b warrants further investigation and may provide insights into plaque development, specifically relating to macrophage infiltration of the vessel wall.

In conclusion, this study provides evidence of the diagnostic potential of TSPO imaging as a means of studying atherosclerotic lesion progression. These results show TSPO expression in a distinct macrophage population that plays an important role in atherosclerotic lesion progression. There is thus a significant rationale for the development and use of radiolabelled TSPO ligands as tools for the detection and quantification of atherosclerotic lesion progression.

## Acknowledgements

This work was supported by an AINSE Research Award to BJC. The authors acknowledge the facilities, scientific and technical assistance of the National Imaging Facility, at the ANSTO-BMC imaging platform, University of Sydney.

